# The Min system disassembles FtsZ foci and inhibits polar peptidoglycan remodeling in *Bacillus subtilis*

**DOI:** 10.1101/872325

**Authors:** Yuanchen Yu, Jinsheng Zhou, Felix Dempwollf, Joshua D. Baker, Daniel B. Kearns, Stephen C. Jacobson

## Abstract

A microfluidic system coupled with fluorescence microscopy is a powerful approach for quantitative analysis of bacterial growth. Here, we measure parameters of growth and dynamic localization of the cell division initiation protein FtsZ in *Bacillus subtilis*. Consistent with previous reports, we find that after division FtsZ rings remain at the cell pole, and FtsZ ring disassembly coincides with rapid Z-ring accumulation at the midcell. In cells mutated for *minD*, however, the polar FtsZ rings persist indefinitely, suggesting that the primary function of the Min system is in Z-ring disassembly. The inability to recycle FtsZ monomers in the *minD* mutant results in maintenance of multiple Z-rings simultaneously, that are restricted by competition for newly synthesized FtsZ. Whereas the parameters of FtsZ dynamics change in the *minD* mutant, the overall cell cycle remains the same, albeit with elongated cells necessary to accumulate a threshold concentration of FtsZ for promoting medial division. Finally, the *minD* mutant characteristically produces minicells composed of polar peptidoglycan shown to be inert for remodeling in the wild type. Polar peptidoglycan, however, loses its inert character in the *minD* mutant suggesting that not only is the Min system important for recycling FtsZ but also may have a secondary role in the regulation of peptidoglycan remodeling.

**IMPORTANCE:** Many bacteria grow and divide by binary fission in which a mothercell divides into two identical daughter cells. To produce two equally sized daughters, the division machinery, guided by FtsZ, must dynamically localize to the midcell each cell cycle. Here, we quantitatively analyze FtsZ dynamics during growth and find that the Min system of *Bacillus subtilis* is essential to disassemble FtsZ rings after division. Moreover, a failure to efficiently recycle FtsZ results in an increase in cell size. Finally, we show that the Min system has an additional role in inhibiting cell wall turnover and contributes to the “inert” property of cell walls at the poles.

## INTRODUCTION

*Bacillus subtilis* is a gram-positive rod-shaped bacterium that grows and divides by a process called binary fission, in which cells increase in mass and divide into two daughters of roughly equal size. During growth, the cell elongates by inserting new peptidoglycan into the lateral cell wall (1). As biomass increases, replication of the chromosome is initiated, and the chromosomes segregate such that the bulk of the nucleoids become evenly spaced within the cytoplasm (2). Cell division is initiated near the geometric midpoint of the cell where peptidoglycan synthesis is reoriented inward towards the cytoplasm to build a septum and complete cytokinesis (3). Medial positioning of cell division ensures that the septum forms between the two nucleoids, guaranteeing each daughter receives one copy of the chromosome. Although the mechanisms governing growth and cell division-site selection are complex, one of the first factors involved in cell division is the protein, FtsZ.

FtsZ is a homolog of eukaryotic tubulin and exists in two different states in the cytoplasm, either as soluble monomers or in long filamentous polymers called protofilaments (4, 5). The two states rapidly interchange as protofilaments dynamically travel by a process called treadmilling in which FtsZ monomers are added to one end and lost from the other (4, 6–11). Treadmilling protofilaments form on the cytoplasmic facing of the membrane and coalesce into a bright focus called the Z-ring at the nascent site of cell division (12–14). Once mature, the Z-ring recruits a transmembrane complex of proteins known as the divisome that synthesizes peptidoglycan on the outside of the cell (15, 16). The Z-ring constricts, either on its own or aided by divisome-directed peptidoglycan synthesis, until the septum is complete, resulting in cytokinesis (17, 18). Thus, FtsZ is both dynamic and seemingly static when concentrated at the site of cell division, and one of the first recognized factors in controlling FtsZ dynamics and localization is the Min system.

The Min system was first discovered in *E. coli* in the form of a mutant that produced minicells at high frequency (19). Minicells are small, metabolically active, spherical bodies that lack DNA and arise when cell division occurs, not at the midcell, but rather near one cell pole. Polar division was attributed to a mislocalization of FtsZ rings and the recruitment of the same machinery that would ordinarily promote medial septation (20). The mutation responsible for minicell formation was mapped to a genetic locus encoding the membrane-associated ATPase MinD and the FtsZ-inhibitor MinC (21, 22). MinD is anchored to the membrane by an amphipathic helix, and MinD recruits and activates MinC by direct interaction (23–30). MinC binds to the C-terminus of FtsZ and destabilizes the FtsZ ring (31–35). In *E. c*oli, the activity of the MinCD complex is dynamically restricted to the polar region by oscillation in which MinCD polar polymerization is antagonized by MinE-mediated depolymerization (36–40). In *B. subtilis* however, the activity of the MinCD complex is statically restricted to membranes with high curvature by MinJ/DivIVA, such that the entire 4-protein complex assembles at the invaginating nascent division plane and remains at the cell poles after division (41–46).

Here, we use fluorescence microscopy and microfluidics to quantitatively measure parameters of *B. subtilis* FtsZ dynamics and cell division under the condition of chemostatic growth for extended periods of time (47–52). The automated poly(dimethylsiloxane) microfluidic system comprises a pneumatically actuated channel array of 600 channels having widths from 1.0 to 1.6 μm and heights of 1.0 μm to actively trap bacteria cells (52). Integrated pumps and valves perform on-chip fluid and cell manipulations that provide dynamic control of cell loading and nutrient flow, and the channel array confines bacterial growth to a single dimension, facilitating high-resolution, time-lapse imaging and tracking of individual cells over multiple generations. In wild type cells, we find that Z-rings persist for a period greater than one cell cycle because Z-rings transiently remain at the cell poles following septum completion. We further show find that the primary function of the Min system is Z-ring disassembly such that in the absence of Min, Z-rings persist longer than the duration of the experiment. Indefinite Z-ring persistence results in cells with multiple Z-rings per compartment, and Z-ring must directly compete for newly synthesized FtsZ monomers. Moreover, we show that *min* mutant cells are elongated because of a failure to recycle monomers, and competition between multiple Z-rings necessitates a larger FtsZ pool. Finally, we provide evidence that the *B. subtilis* Min system also inhibits cell wall turnover, particularly at the poles of the cell, and is a contributing factor to reports of inter-polar peptidoglycan.

## MATERIALS AND METHODS

### Strains and growth conditions

*B. subtilis* strains were grown in lysogeny broth (LB) (10 g tryptone, 5 g yeast extract, 10 g NaCl per L) or on LB plates fortified with 1.5% Bacto agar at 37°C.

### Microfluidic system

The microfluidic device was fabricated through a combination of electron-beam lithography, contact photolithography, and polymer casting (52). Briefly, the microfluidic device is comprised of fluid and control layers both cast in poly(dimethylsiloxane) (PDMS) and a glass coverslip. The fluid layer lies between the control layer and glass coverslip and contains the microchannels and channel array to trap the bacteria. Media and cells are pumped through the microfluidic channels by on-chip peristaltic pumps and valves that are controlled pneumatically through the top control layer. Each pneumatic valve is controlled by software to apply either vacuum (0.3 bar) or pressure (1.3 bar) to open or close individual valves, respectively. Device fabrication design details are included in Figure S1.

### On-device cell culture

Prior to loading cells into the microfluidic device, the fluidic channels were coated with 1% bovine serum albumin (BSA) for 1 h to act as a passivation layer. Then, all the channels were filled with 1 mM IPTG, 0.1% BSA in Luria-Bertani (LB) media (10 g tryptone, 5 g yeast extract, 10 g NaCl per L). A saturated culture of cells (∼25 μL) was added through the cell reservoir and pumped into the cell-trapping region. During cell loading, vacuum was applied to the control layer above to open the membrane region. After a sufficient number of cells were pumped underneath the channel array, positive pressure was applied to trap individual cells in those channels. Media was pumped through the microchannels to flush away excess cells. After excess cells were pumped away, media was continuously flowed through the microchannels by gravity flow during the entire experiment.

### Time-lapse image acquisition

Steady state cell growth was monitored from 3 to 21 h post inoculation. Fluorescence microscopy was performed on a Nikon Eclipse Ti-E microscope and an Olympus IX83 microscope. The Nikon Eclipse Ti-E microscope was equipped with a 100x Plan Apo lambda, phase contrast, 1.45 N.A., oil immersion objective and a Photometrics Prime95B sCMOS camera with Nikon Elements software (Nikon, Inc.). Fluorescence signals from mCherry and mNeongreen were captured from a Lumencor SpectraX light engine with matched mCherry and YFP filter sets, respectively, from Chroma. The Olympus IX83 microscope was equipped with an Olympus UApo N 100x/1.49 Oil objective and a Hamamatsu EM-CCD Digital Camera operated with MetaMorph Advanced software. Fluorescence signals from mCherry, mNeongreen and BADA were captured from an Olympus U-HGLGPS fluorescence light source with matched TRITC and GFP filters, respectively, from Semrock. Images were captured from at least eight fields of view at a 2 min interval. The channel array was maintained at 37°C with an objective warmer. For all direct comparisons, the same microscope and settings were used.

### Data analysis

A period of adaptation following exposure to illumination was observed; thus, data analysis was restricted to periods of steady state. Cell identification and tracking were analyzed by a series of MATLAB programs (The MathWorks, Inc.) (52). The program extracted fluorescence intensity along a line profile down the longitudinal center of each sub-micron channel. The cytoplasmic mCherry line profile showed a flat topped peak on the line where a cell was located, and a local 20% decrease in fluorescence intensity was used to identify cell boundaries after division. Division events were conservatively measured as the time at which one cell became two according to the decrease in fluorescence intensity as described above. Moreover, cell bodies were tracked from frame to frame in order to construct lineages of cell division. The mNeongreen-FtsZ signal was similarly tracked and measured along the length of the cell. The FtsZ line profile was normalized by cell body intensity in order to minimize intensity differences among frames and different fields of view.

### Fluorescent D-amino acid labeling

The fluorescent D-amino acid, BADA, was supplied by VanNieuwenhze and Brun labs. To create BADA, 3-amino-D-alanine was conjugated to BODIPY-FL (53, 54). Stock solutions of BADA (100 mM) were prepared in dimethylsulfoxide (DMSO) as they were poorly soluble in water. The stock solutions were then diluted with LB media to 1 mM BADA with < 2% (vol/vol) DMSO left. To label the cells 3 h post inoculation, the BADA solution (1 mM) was continually pumped through the channel array for 4 min. Excess dye was washed away by pumping LB media through the channel array for 4 min. Fluorescent images were captured at a 2 min interval from at least eight fields of view.

## RESULTS

### The Min system is required for FtsZ ring disassembly

Quantitative microscopic analysis of cell growth and division on agarose pads is restricted by the limited number of generations that can be observed under batch conditions. To circumvent this problem, a microfluidic-based approach was undertaken to monitor steady-state chemostatic growth of *B. subtilis* over many generations (**Fig S1**). *B. subtilis* divides by septation (or plate formation) in which a division septum is formed first, and remodeling of the septal peptidoglycan occurs as a separate step afterwards that leads to indentation and cell separation (55–57). Thus, cell division events were conservatively defined as a spatial decrease in constitutively-expressed cytoplasmic mCherry fluorescent signal that would indicate cellular indentation (Fig 1A). Images were captured every two minutes, and the fluorescence intensity was measured along the length of the microfluidic channel. After inoculation into the microfluidic device, a period of roughly three hours elapsed during which cells appeared to adjust to the growth conditions, and steady-state growth was maintained and monitored over the next 21 hours (**Movie S1**). Microscopic analysis of the rate of septum formation indicated that wild type cells grew with a cell cycle of 39 ± 12 min (Fig 2A).

**Figure 1:**
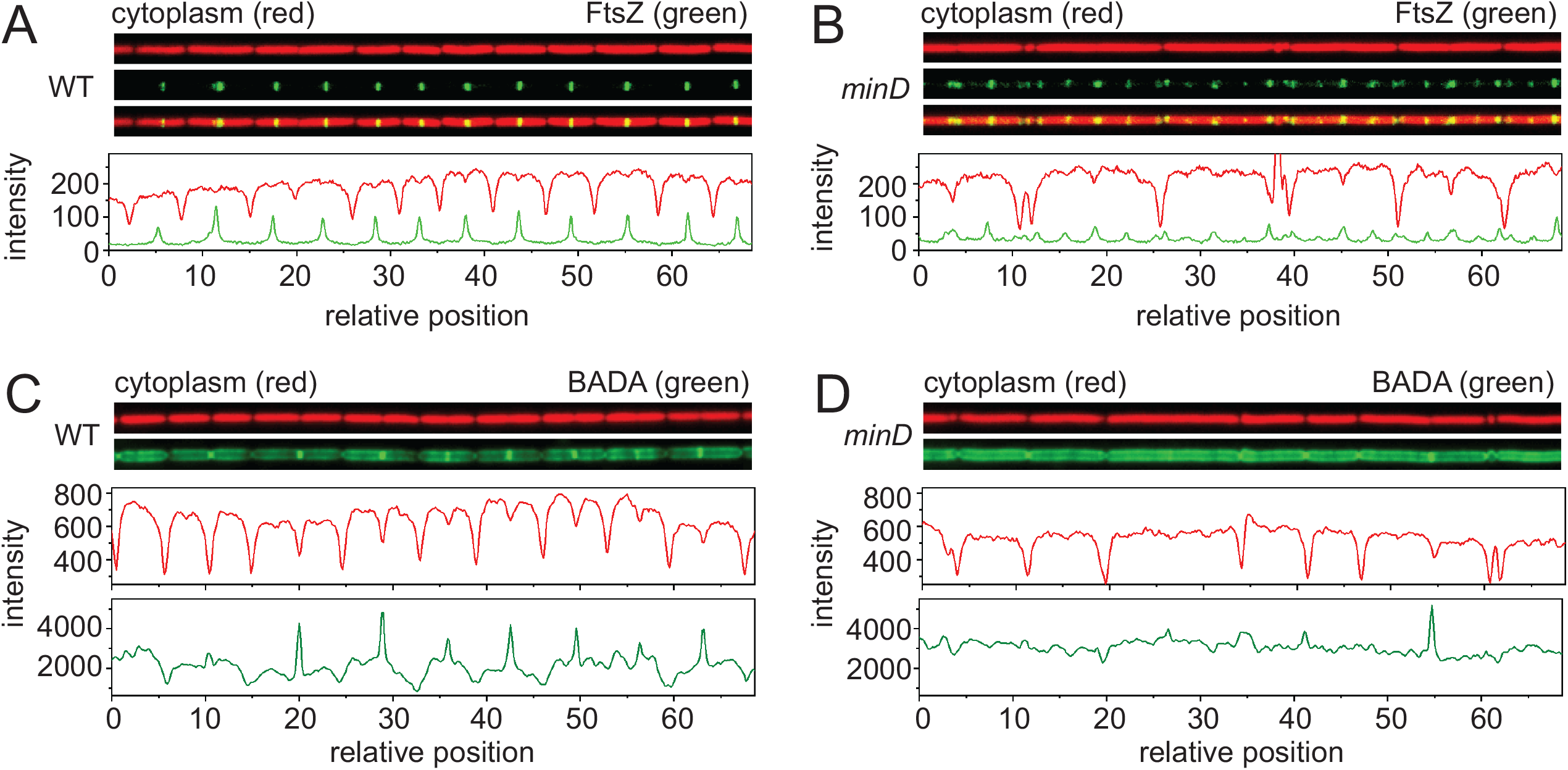
Microfluidic analysis of growth and division in wild type and *min* mutants. Snapshot fluorescent microscopy of a microfluidic channel for wild type (A,C) and a *minD* mutant (B,D) growing at steady state. A and B) Fluorescence microscopy of cells in a microfluidic channel expressing cytoplasmic mCherry protein false colored red (top), mNeongreen-FtsZ false colored green (middle), and an overlay of the two colors (bottom). Graphs are a quantitative analysis of mCherry fluorescence intensity (red line) and mNeongreen fluorescence intensity (green line) to match the fluorescence microscopy images immediately above. All images are reproduced at the same magnification. C and D) Fluorescence microscopy of a microfluidic channel for wild type (A,C) and a *minD* mutant (B,D) growing at steady state. Cytoplasmic mCherry protein false colored red (top), and peptidoglycan stained with BADA false colored green (bottom). Graphs are a quantitative analysis of mCherry fluorescence intensity (red) and BADA fluorescence intensity (green) to match the fluorescence microscopy images immediately above. All images are reproduced at the same magnification. Wild type (DK5133) and a *minD* mutant (DK5155) were used to generate all of the data in this figure.

**Figure 2:**
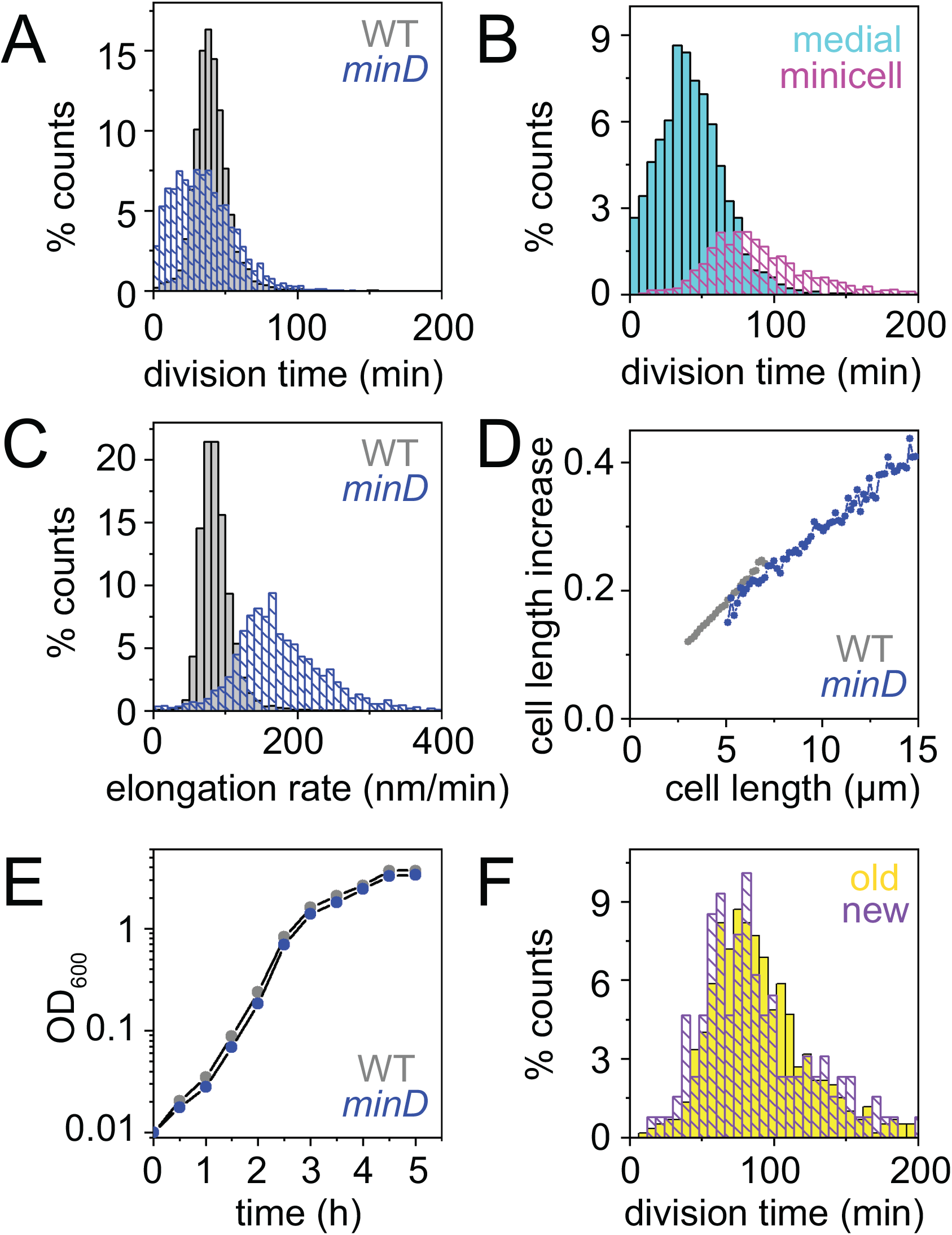
Cells mutated for the *min* system divide faster than wild type. A) A histogram of the division time of individual cells of wild type (gray) and a *minD* mutant (blue) measured by microscopic analysis. Division events were defined by a local 20% decrease in mCherry (cytoplasmic) fluorescence intensity below a threshold value. More than 3000 division events were counted per dataset. Minicells were excluded from the division time analysis as once they are formed, minicells never divide. B) A histogram of division time of individual cells of the *minD* mutant using the dataset in panel A to separately determine the time elapsed between medial and minicell divisions. The time between medial division events (cyan) was determined as those events that gave rise to two separate cells with chromosomes. The time between minicell divisions (magenta) was determined to be the time between the formation of a cell pole and the formation of a division plane at that pole to give rise to a minicell. C) Cell elongation rates were measured as the rate at which the cell poles moved apart from one another in wild type (gray) and a *minD* mutant (blue). The growth of over 2500 cells was measured for each strain. Minicells were excluded from the elongation rate analysis as once they formed, minicells do not elongate. D) Data from panel 2C was replotted as the instantaneous increase in cell length per total length of the cell observed. E) Growth curve of wild type (gray) and a minD mutant (blue) growing in highly agitated LB broth at 37°C and optical density was measured with a spectrophotometer at 600 nm wavelength. Wild type (DK5133) and a *minD* mutant (DK5155) were used to generate all of the data in this figure. F) Frequency histogram of division time that gives rise to minicells at either the old cell pole (yellow) or the new cell pole (violet). Old cell poles were defined by the pole that last experienced a polar division event. New cell poles were defined by the pole that had not previously experienced a polar division event.

Cell division is mediated by dynamic localization of the division initiation protein, FtsZ (8, 9, 12, 14). FtsZ dynamics were monitored by fluorescence microscopy in a strain encoding an N-terminal fluorescent fusion of mNeongreen introduced at the native site in the chromosome (9). Images were captured every two minutes, and the fluorescence intensity magnitude of FtsZ was measured as a snapshot in the context of a fluorescent mCherry cytoplasmic signal (Fig 1A). During steady state growth, FtsZ appeared as a faint uniform cytoplasmic haze with bands of enhanced fluorescence intensity, and a kymograph was generated to track FtsZ dynamics in temporal relation to the cell body (Fig 3A, Fig 1A**, Movie S2**). The FtsZ-ring appearance period, defined as the time between the formation of one Z-ring and the formation of another, was found to be similar to that of the cell division period (Fig 4**, Fig S2A**). The Z-ring persistence period, defined as the time between appearance and disappearance of a single focus, was longer than the average period of cell division (Fig 4**, Fig S2B**) likely because FtsZ has been observed to remain transiently at the cell pole after cytokinesis (13, 46, 58). Consistent with previous observations, many cells exhibited a characteristic peak of FtsZ fluorescence intensity near midcell to mediate division, but some cells instead exhibited peak fluorescence at the cell pole after division was complete (Fig 5A). We conclude that FtsZ remains polarly localized after cytokinesis.

**Figure 3.**
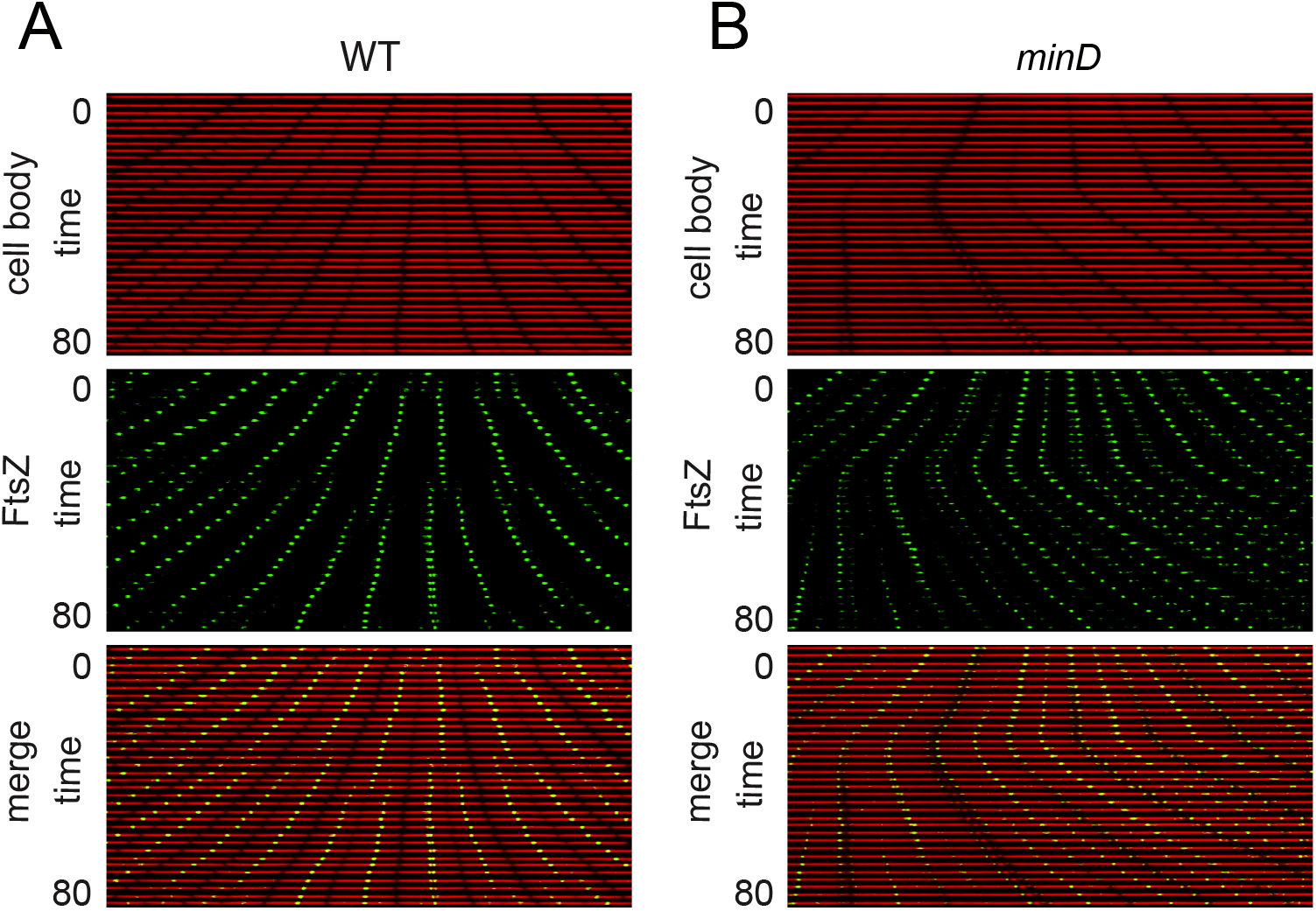
FtsZ foci remain at the poles indefinitely in the absence of Min. Kymograph analysis of wild type (A) and a *minD* mutant (B) of cytoplasmic mCherry signal false colored red (top) and mNeongreen-FtsZ intensity false colored green (middle), and an overlay of the red and green channels (bottom). In each panel a single microfluidic channel was followed in a series of stacked snapshots taken at 2 minute intervals to assemble the kymograph. All images reproduced at the same magnification. Wild type (DK5133) and a *minD* mutant (DK5155) were used to generate all of the data in this figure.

**Figure 4:**
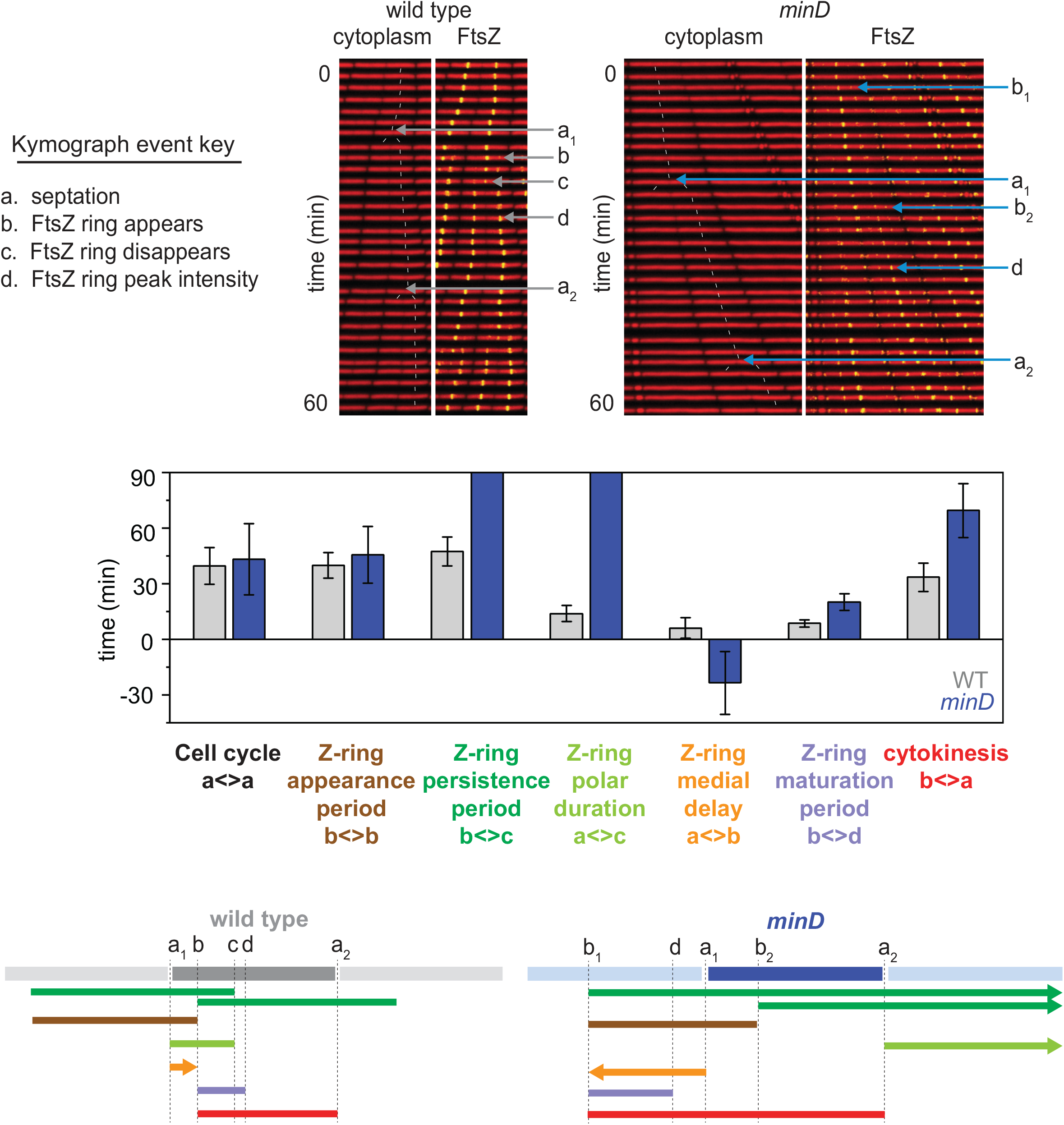
The cytokinetic period is longer than a cell cycle in a *minD* mutant. Top) Sample kymograph analysis of a wild type (A) and a *minD* mutant (B) microfluidic channel in which cytoplasmic mCherry signal is false colored red (left) and overlayed with mNeongreen-FtsZ that is false colored green (right). Events necessary for defining division paramaters are indicated and labeled a, septation; b, detection of a nascent Z-ring; c, disappearance of a Z-ring; d, FtsZ peak intensity achieved. Thin white lines are include to indicate cell tracking and lineage analysis. Middle) Graphs of 100 manually tracked wild type (gray) and minD (blue) cell cycles presented as bars of average values and whiskers of standard deviation for the following parameters: “Cell cycle” is the time between septation events (between consecutive “a” events); Z-ring appearance period is the time between the formation of one Z-ring and another (between consecutive “b” events); “Z-ring persistence period” is the time between the formation of a Z-ring and the disappearance of that Z-ring (between consecutive “b” and “c” events); “Z-ring polar duration” is the time between a septation event and the disappearance of the Z-ring resulting from that septation events (between consecutive “a” and “c” events); “Z-ring medial delay” is the time between a septation event and the formation of a Z-ring that will eventually give rise to the next medial division event (between an “a” event and a “b” event that will give rise to the next round of septation); “Z-ring maturation period” is the time between Z-ring formation and when that Z-ring achieves peak local intensity (between consecutive “a” and “d” events); and the “cytokinetic period” is the time between Z-ring formation and septation directed by that Z-ring (between a “b” event and the “a” event that is caused by that particular Z-ring). The raw data histograms for each bar are presented in Fig S2. Bottom) Timeline representations of the various events indicated in the bar graph depicted in cartoon form, color coded to match the indicated parameter of like color above, and annotated with relevant events marked by the defining letters.

**Figure 5:**
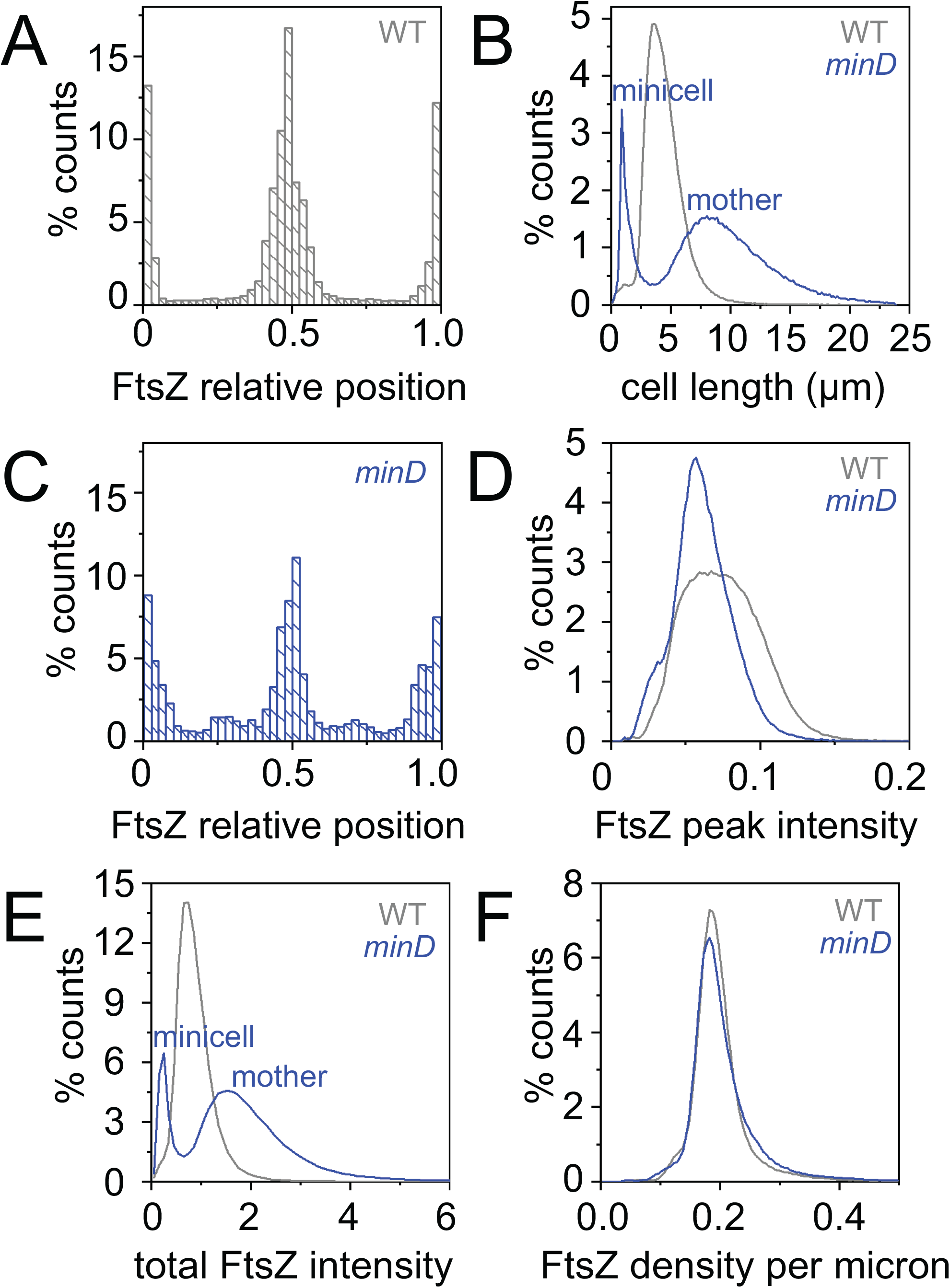
FtsZ density is constant in both the wild type and the *minD* mutant. A) A frequency histogram of the location of snapshot peak FtsZ intensity plotted relative to total cell length for wild type. The poles of the cell have relative position values of 0 and 1.0 whereas the midcell has a value of 0.5. B) A frequency histogram of cell length distribution of wild type (gray) and a *minD* mutant (blue). The *minD* mutant has two peaks, a shorter peak corresponding to “minicells” and a longer peak corresponding to “mother” cells that have chromosomes and are capable of division. C) A frequency histogram of the location of snapshot peak FtsZ intensity plotted relative to total cell length for *minD*. D) A frequency histogram of peak FtsZ fluorescence intensity per cell. For each frame, a line scan through the longitudinal axis of the cell determined the location of peak fluorescence intensity and peak fluorescence magnitude was recorded. Wild type distribution (gray); *minD* distribution (blue). E) A frequency histogram of total FtsZ fluorescence intensity per cell was measured by integrating the area under the line scans from panel D. F) A frequency histogram of total fluorescence intensity was divided by cell length for each individual. Measurements were taken for 9000 cells of growing wild type (DK5133) amounting to over 30,000 measurements, and 3000 cells of growing minD (DK5155) for over 20,000 measurements to generate the data in this figure.

To further explore Z-ring dynamics in the wild type, 100 cells were chosen at random, and a variety of parameters were manually measured. The Z-ring polar duration, defined as the time between septation and the disappearance of the Z-ring (Fig 4**, Fig S2C**) was longer than, and overlapped with, the Z-ring medial delay (Fig 4**, Fig S2D**), defined as the time between septation and the formation of a new Z-ring. Thus, wild type cells transiently experienced multiple FtsZ rings per compartment. Moreover, dissolution of the polar Z-ring coincided with the Z-ring maturation period (Fig 4**, Fig S2E**), defined as the time from first appearance of the Z-ring until maximum Z-ring fluorescence intensity, as FtsZ subunits were redistributed from the pole to the midcell. Finally, the cytokinetic period, defined as the time between Z-ring appearance and cell division was approximately 33 ± 8 min (Fig 4**, Fig S2F)**, which when added to the Z-ring medial delay, ultimately produced a value similar to the cell cycle period (39 ± 12 min) (Fig 4). We conclude that the dynamic parameters of FtsZ are consistent with a regular cell division cycle despite the transient localization of the Z-ring at the poles.

One mechanism that governs FtsZ localization is the Min system (20, 21, 59). To explore the consequences of disruption of the Min system quantitatively, a mutation was introduced in the gene encoding MinD, the membrane localized activator of the FtsZ-inhibitor protein MinC, and the *minD* mutant was monitored during growth in microfluidic channels (Fig 1B). Consistent with a *min* phenotype, the *minD* mutant produced two different cell types: cells that were longer than wild type and very short minicells (Fig 5B**; Movie S3**) (19, 21, 22, 60). Cells mutated for MinD produced multiple FtsZ foci (61) (Fig 1B**; Movie S4**) and kymograph analysis indicated that the FtsZ-ring persistence period was indefinite such that once formed, the focus did not disappear during the timecourse of observation (Fig 3B). Moreover, a greater proportion of *minD* mutant cells exhibited peak FtsZ fluorescence intensity at the poles (Fig 5C). We conclude that when a cell divides, the FtsZ focus is split into each daughter cell. In the wild type, the polar FtsZ ring is transient due to antagonism by Min, but in the absence of Min, the ring persists. Our data support models in which the primary function of the Min system during division is to promote FtsZ ring disassembly rather than preventing its formation (23, 46, 62).

### The Min system maintains cell size by recycling FtsZ

Mother cells of the *minD* mutant (8.8 ± 3.4 μm, not including minicells) were on average twice as long as the wild type (4.5 ± 1.7 μm) (Fig 5B), but the reason *min* mutants were elongated was unclear. One early model suggested that *min* mutants were longer because divisions that produced minicells came at the expense of medial divisions (60, 63). Microscopic analysis, however, indicated that the division time in the *minD* mutant occurred slightly more rapidly than the wild type (Fig 2A). By considering divisions that gave rise to different cell types separately, one quarter of all division events in *minD* gave rise to minicells, and three quarters of division events occurred along the midcell to produce two viable daughters with chromosomes (Fig 2B). Thus, midcell divisions occurred at roughly the same average rate as the wild type albeit with a higher standard deviation (Fig 2B). The wide variance was due to the occasional longer-than-average division times that gave rise to very long cells which then experienced shorter-than-average division times with multiple division events per compartment that could occur simultaneously or slightly offset from one another (Fig 6 ) (19, 64). We conclude that *min* mutants experience medial division at approximately the same rate as the wild type, and thus, *min* cells are not elongated because polar septation comes at the expense of medial division.

**Figure 6.**
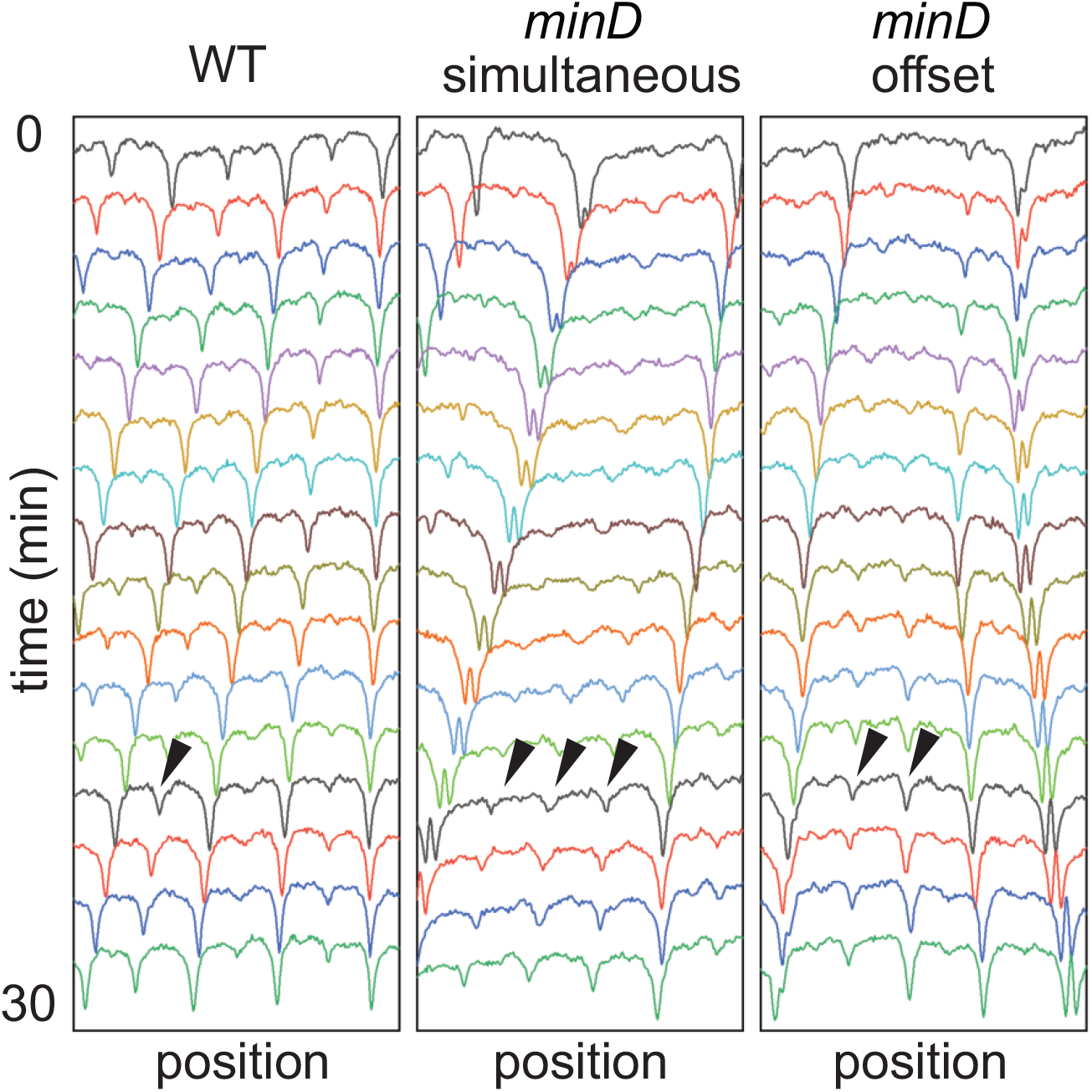
Some cells of a *minD* mutant experience multiple simultaneous division. Kymograph analysis of cytoplasmic mCherry fluorescence intensity of single channels of a wild type (left) or *minD* (middle and right) mutant. Wild type cells experience regular division once per cell cycle (caret, left panel) but occasionally multiple simultaneous (carets middle panel) or slightly offset (carets right panel) division events were observed in the *minD* mutant. The presented kymographs of individual wells were representative of general trends taken from the image stacks generated in Figure 1. Wild type (DK5133) and a *minD* mutant (DK5155) were used to generate all of the data in this figure.

Another factor that could contribute to cell length is the rate of cell elongation, as a faster elongation rate relative to the division rate could give rise to longer cells. By measuring the rate at which the cell poles moved away from one another, the *minD* mutant appeared to elongate more rapidly than the wild type (Fig 2C). Elongation, however, occurs by the lateral synthesis of cell wall material distributed along the length of the rod (65–67), and thus, the *minD* mutant might appear to elongate more rapidly simply because longer cells have more positions in between the poles in which to insert new cell wall material per unit time. However, the instantaneous elongation rate, defined as the rate of elongation divided by the length of the cell, was similar for both the wild type and *minD* mutant (Fig 2D). Moreover, cells of the wild type and the *minD* mutant accumulated biomass at the same rate as both had indistinguishable growth curves by optical density measurement in broth culture (Fig 2E). We conclude the cells of the *minD* mutant elongate at the same rate as the wild type, and thus, the elongation rate cannot explain the presence of elongated cells.

Another possible explanation for the increased cell size of *min* mutants is a reduced rate at which FtsZ monomers are added to form FtsZ rings. “Adder” hypotheses for cell growth predict that cell size is dictated by the accumulation of a critical threshold concentration of a particular cell component, in this case, FtsZ (60, 68–70). One requirement for the adder hypothesis is that the rate of FtsZ accumulation must be constant. With respect to FtsZ accumulation, the magnitude of peak FtsZ fluorescence intensity was higher in the wild type than the *minD* mutant (Fig 5D), but the *minD* mutant exhibited higher total fluorescence intensity per cell (Fig 5E). The higher total fluorescence intensity per cell may be due to the fact that the *minD* cells are longer and maintain multiple Z-rings per compartment. Indeed, when fluorescence intensity was divided by cell length, FtsZ density was found to be nearly identical in both *minD* and the wild type. Thus, the rate of FtsZ synthesis was constant, and FtsZ accumulation was proportional to cell size (Fig 5F). We conclude the *minD* mutant cells are elongated because old Z-rings are not recycled, and the cell needs more time and, thus, more biomass to accumulate FtsZ sufficient to form and mature a new Z-ring. We further infer that Z-ring formation and, subsequently, cell division in *minD* directly depend on the rate of *de novo* FtsZ synthesis.

If FtsZ density accumulated at a rate proportional to cell length (Fig 5F), and the *minD* mutant constitutively maintained multiple Z rings, new FtsZ monomers synthesized during growth were likely divided between multiple foci, effectively reducing the rate of local FtsZ accumulation that gives rise to cytokinesis. Indeed, the nascent Z-rings of the *minD* mutant exhibited a 2-3 fold increase in maturation time (Fig 4**, Fig S2E**), likely because the old rings were not dissolved, and each Z-ring independently competed for newly synthesized subunits. We note that the cytokinetic period between FtsZ formation and septation was 2-3 fold longer (Fig 4**, Fig S2F**), suggesting that regeneration of cell division machinery was also delayed. Paradoxically, the cytokinetic period was substantially longer than the cell cycle (Fig 4). Each daughter cell of the *minD* mutant, however, was born with at least three persistent Z-rings, one at each pole and one at the future midcell, and additional Z-rings formed at the one-quarter and three-quarter position as the cells grew (Fig 5C). Thus, division at the most central Z-ring allowed daughter cells to be born with a preformed medial ring that would eventually drive septation, and as a result the medial Z-ring delay time for *minD* had a negative value (Fig 4**, Fig S2D)**. Despite aberrations in Z-ring parameters, the minD mutant still maintained a similar cell cycle time as the wild type, correlated not with the cytokinetic period but rather with the rate of FtsZ ring appearance (Fig 4**, Fig S2A**). We conclude that the Min system disassembles polar Z-rings to recycle and redistribute monomer units, thereby promoting rapid, singular FtsZ accumulation and maturation at a medial site for the proper maintenance of both the cytokinetic period and cell length (Fig 4).

### The Min system inhibits peptidoglycan turnover, especially at the cell poles

Mutants defective in the Min system not only produce longer cells but also produce minicells by division at the cell poles. Peptidoglycan at the cell poles is traditionally considered to be “inert” such that once synthesized, it experiences little *de novo* synthesis and turnover (67, 71–73). To study how minicells formed, cells in the microfluidic device were presented with a sub-generational (4 min) impulse of fluorescent D-amino acids that can be incorporated into peptidoglycan during either synthesis or remodeling (53, 54). In wild type, fluorescent peptidoglycan signal was incorporated along the central part of the cell body with peak fluorescence at the nascent division planes (Fig 1C). Moreover, the troughs of peptidoglycan fluorescence intensity coincided with troughs of cytoplasmic fluorescence intensity consistent with the idea that polar peptidoglycan was less dynamic than other positions (Fig 1C). Peptidoglycan staining of the *minD* mutant, however, appeared more intense and more uniform than the wild type (Fig 1D). In both wild type and the *minD* mutant, peptidoglycan staining was proportional to cell length, but the *minD* mutant accumulated more stain per unit length (Fig 7A). Also, unlike the wild type, troughs in cytoplasmic staining intensity in the *minD* mutant did not correspond to decreases in peptidoglycan staining at the cell poles (Fig 1D). We conclude that the Min system inhibits peptidoglycan synthesis/remodeling and is an important factor in making the polar peptidoglycan appear “inert”.

**Figure 7.**
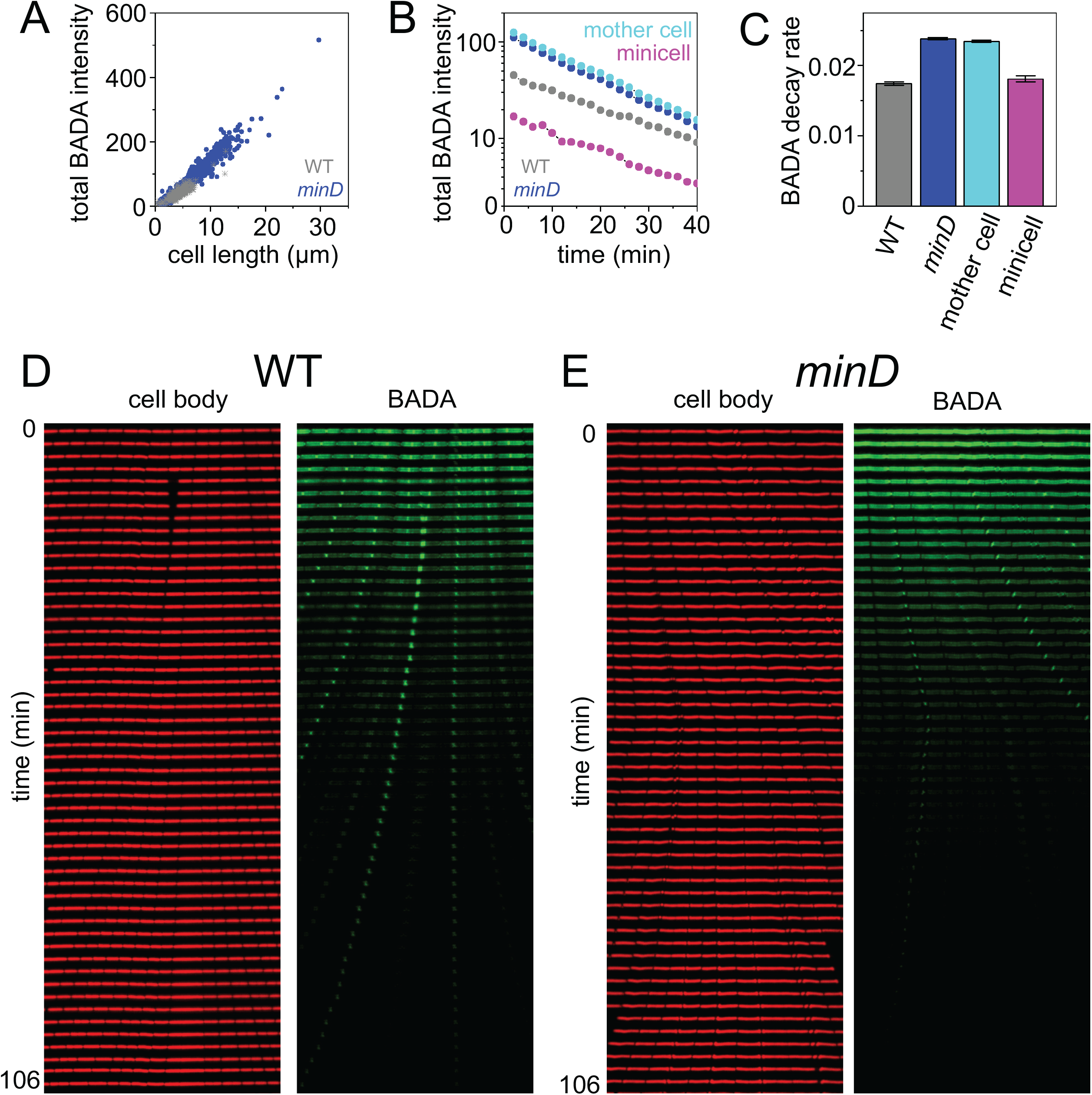
*minD* mutants lose BADA staining fluorescence intensity faster than the wild type. Cells were grown in the presence of the peptidoglycan synthesis/remodeling indicator stain BADA for 4 minutes, the stain was washed out of the microfluidic device for 8 minutes and then fluorescence intensity was tracked in time. A) A graph of the total BADA intensity per cell per the length of the cell measured in the first frame of the experiment. Each dot represents an individual cell of wild type (gray) and *minD* mutant (blue). B) A graph of the total BADA intensity per cell per unit time after washout of fluorescent D-amino acids. Gray, wild type; Blue; *minD* mutant, Cyan, mother cells of the *minD* mutant with chromosomes, and magenta, minicells of the *minD* mutant. C) A graph of the rate of decrease in BADA intensity loss as measured by the slope of the lines in Figure 7B. D) A representative kymograph of wild type after BADA washout. E) A representative kymograph of *minD* after BADA washout. The wild type (DK4393) and *minD* mutant (DK4407) strains were used for all panels in this figure. Over 500 measurements were taken for each strain.

We next wanted to determine how the *minD* mutant appeared to accumulate more fluorescent peptidoglycan signal. As above, cells were stained with a 4-min impulse of fluorescent D-amino acids after which, the fluorescent D-amino acids were replaced with unlabeled D-amino acids in the growth medium such that the fluorescence signal would be gradually lost during growth (**Movie S5,S6**). When the amount of fluorescence signal per cell was projected per unit time, *minD* mutant cells with chromosomes experienced a greater overall rate of fluorescence decay than the wild type or *minD* mutant minicells (Fig 7 B,C, **Fig S3 A,B**). Consistent with inert polar peptidoglycan in the wild type, the most intense staining occurred at the nascent division plane, which upon becoming a cell pole remained fluorescent for an extended period of time (Fig 7D). The *minD* mutant, however, did not exhibit intense staining at the division plane, and polar staining persisted primarily in minicells (Fig 7E). We conclude that the Min system, at least in *B. subtilis*, appears to restrict peptidoglycan remodeling throughout the cell but has the greatest effect at the cell poles.

Minicells are thought to be physiologically similar to wild type cells due to inheritance of cytoplasmic and membrane proteins, but they lack chromosomes and do not grow. One reason minicells might not grow is that they were thought to be surrounded exclusively by inert polar peptidoglycan, but poles of the *minD* mutant appeared much less inert than the wild type (Fig 1D). To further explore the dynamics of polar peptidoglycan observed in the *minD* mutant, the mutant was stained for a longer period of time (20 min). Prolonged exposure of the *minD* mutant to fluorescent D-amino acids resulted in staining of minicell peptidoglycan but only in the most recently formed minicells (Fig 8). In each case the entire circumference of the minicell was stained uniformly indicating that old cell pole peptidoglycan was being remodeled at a rate equivalent to the synthesis of the nascent plane (Fig 8). Little to no fluorescence was observed in older minicells suggesting that minicells rapidly lost their ability to remodel peptidoglycan (Fig 8). We conclude that minicells transiently retain the ability to remodel peptidoglycan, and that the rapid loss of remodeling capacity may be responsible for the inability of minicells to grow. Combined, our microfluidic analysis indicates that the Min system in *B. subtilis* is multifunctional: Min not only disassembles polar FtsZ-rings but also restricts polar peptidoglycan remodeling.

**Figure 8.**
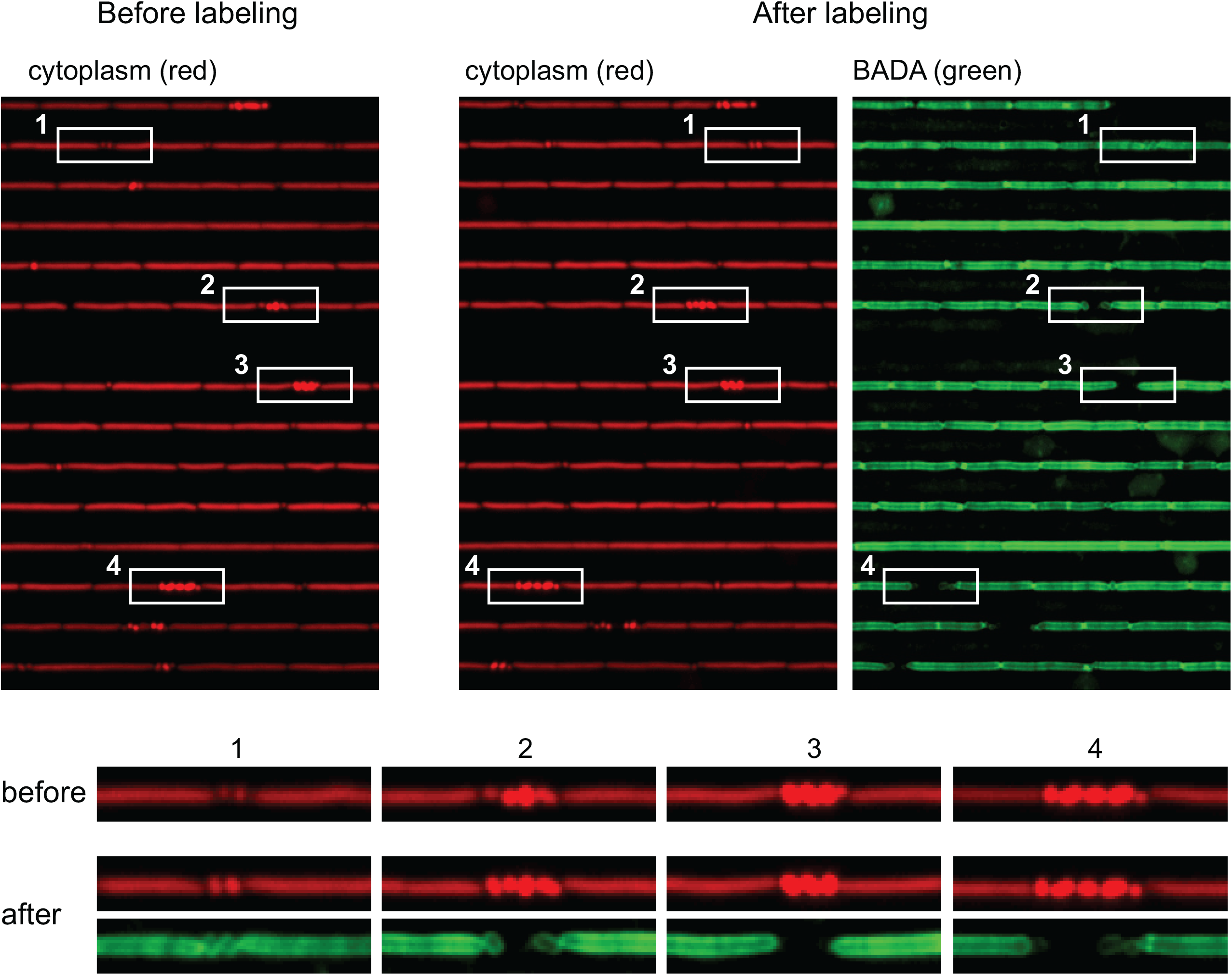
Recently formed minicells are proficient for remodeling of the polar peptidoglycan. A) Microfluidic analysis of a *minD* (DK4407) mutant expressing cytoplasmic mCherry false colored red (left) and stained for 20 minutes with the fluorescent D-amino acid BADA false colored green (right). Individual areas of the channels are highlighted by white boxes and numbered. Below are enlarged images of the boxes with the corresponding number to increase detail.

**Table 1:**
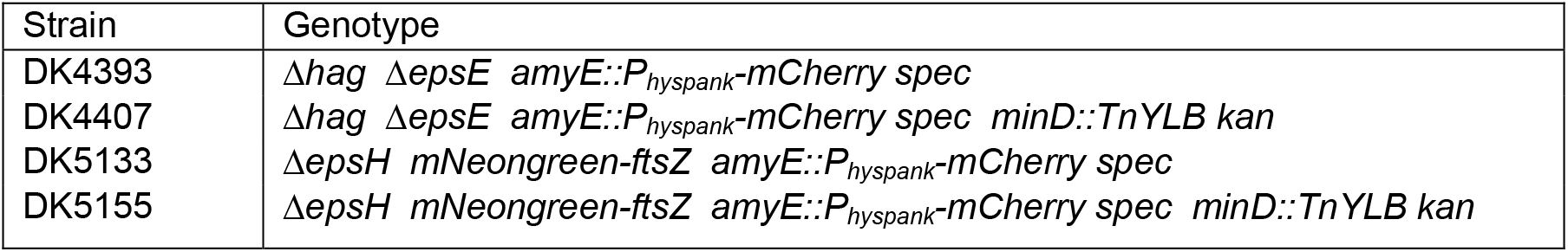
Strains.

## DISCUSSION

Binary fission of rod-shaped bacteria is considered one of the simplest forms of cell division, but it is nonetheless complex. Cells elongate by directing envelope synthesis parallel to the long axis of the cell but must periodically reorient envelope synthesis perpendicular to the long axis to promote cytokinesis. FtsZ forms a ring-like scaffold to recruit the cytokinetic complex for perpendicular peptidoglycan synthesis, and the Min system is thought to help guide FtsZ localization. Here, we quantitatively measure growth and FtsZ dynamics in wild type and a *minD* mutant of *B. subtilis* with a microfluidic device and fluorescence microscopy. We find that the primary function of the Min system is to disassemble FtsZ rings and recycle FtsZ monomers. A failure to recycle FtsZ monomers has a number of consequences on FtsZ dynamics, including the constitutive maintenance of multiple Z-rings per compartment and a longer cytokinetic period. Despite alternations in FtsZ dynamics, however, we find that the *minD* cell cycle is nonetheless much like wild type cell cycle. Moreover, the *min* mutants are so named because they produce small anucleate minicells made exclusively of polar peptidoglycan thought to be relatively inert. Here, we provide evidence that the Min system is responsible for inhibiting polar peptidoglycan turnover and that minicells transiently retain peptidoglycan remodeling capacity.

The function of the Min system is often described as promoting medial Z-ring formation by preventing FtsZ-ring formation at the poles, but our data, along with others, suggest otherwise, at least in *B. subtilis*. Rather than preventing Z-ring assembly, Min is required for FtsZ ring disassembly (23, 30, 35, 46, 62, 74). Evidence that Min does not occlude polar Z-rings comes from spore outgrowth experiments which indicate that FtsZ preferentially localizes to the midcell even when *minD* is mutated, and thus, the *min* phenotype does not manifest in the first generation (75, 76). Instead, FtsZ transiently remains at the cell pole after division and is redistributed to the midcell by the action of Min, such that in the absence of the Min system, FtsZ rings are divided into each new daughter but never dissolve. A Z-ring disassembly model is also consistent with the fact that the Min system co-localizes with nascent septa behind the constricting ring and only encounters FtsZ after septation has completed (77). A disassembly function is also supported by genetic evidence showing that mutants with enhanced Z-ring stability are resistant to the effects of MinCD and mutants with reduced Z-ring stability have enhanced sensitivity (33, 78, 79). That FtsZ may require a specific disassembly factor may also be consistent with the division of *Streptococcus pneumoniae*, which naturally lacks the Min system and produces Z-foci that are not depolymerized. These Z-foci, instead, continually treadmill from the old division site to the new (73).

The inability to disassemble Z-rings and recycle FtsZ monomers also explains cell length defects associated with *min* mutants. *min* mutants have long been observed to not only produce minciells but also have mother cells that are longer than the wild type (19, 21, 22, 60). Here, we show that *min* mutants were not elongated due to an extended division time or increased elongation rate but rather due to the dynamics of FtsZ accumulation. In the absence of Min, the persistence of the Z-ring becomes infinite, and cells maintain multiple Z-rings per compartment. Because the density of FtsZ is constant, each Z-ring is forced to compete for newly synthesized FtsZ monomers. Thus, the 2-3 fold increase in cell length was correlated with a 2-3 fold increase in Z-ring maturation time. Our results are consistent with the “adder” hypothesis for growth and cell length control, as *min* mutant cells must become longer to achieve a critical threshold of FtsZ to promote cell division (60, 68–70). Additionally, we observed that the *minD* mutant had a 2-3 fold increase in the cytokinetic period. We conclude that not only was there a failure to recycle FtsZ, but cell division machinery was co-sequestered at latent Z-rings and also required *de novo* synthesis for regeneration (74).

Min-mediated disassembly of the latent polar Z-ring coincided with rapid accumulation of a medial Z-ring in the wild type cell cycle. Despite defects in FtsZ dynamics, the *minD* mutant did not suffer a defect in either growth rate or division time, suggesting that the cell cycle was robust and depended on another factor. In the absence of Min, the cytokinetic period exceeded the cell cycle time, Z-ring formation that would give rise to the future division was initiated in the preceding generation. Thus, much like how multifork replication allows daughters to inherit partially replicated chromosomes and grow faster than the replication period, daughter cells inherit a mature Z-ring to complete the cell cycle on time and assemble two new Z-rings for the next generation. Ultimately, the cell cycle of the *min* mutant is governed by the rate of Z-ring appearance, but what governs the Z-ring appearance rate is unclear. One likely regulator of Z-rings is the chromosome because Z-rings are prevented from forming over the mass of the genetic material by nucleoid occlusion (80, 81). We infer that the cell cycle is preserved because the regular period of chromosome replication and segregation dictates the rate at which Z-rings appear, and we note that the Z-ring appearance period is similar to the replication period reported in *B. subtilis* (58, 82, 83).

The failure to disassemble Z-rings in the absence of Min leaves behind a preformed polar Z-ring that can give rise to polar cytokinesis and results in the classic phenotype of minicell division. We note, however, that despite the fact that each daughter inherits three Z-rings, two at the pole and one at the midcell, and each Z-ring sequesters division machinery, the division events that gave rise to minicells were nonetheless rare. Competition between the Z-rings for newly synthesized FtsZ monomers and divisome components is unequal, such that the medial Z-ring is stochastically favored. We suspect that medial rings are favored because they are flanked on either side by two chromosomes, each expressing divisome components, and polar Z-rings are disfavored by diffusion being proximal to only one chromosome. Consistent with a positional bias in diffusion-and-capture, medial divisions occur at approximately twice the frequency of minicell divisions, and we found that minicell formation at the new and old pole was equally probable (Fig 2F). Regardless on which side of the cell polar-division occurs, the minicell compartment is reduced to a sphere in which half of the peptidoglycan comes from one pole and half comes from the nascent septation event.

Polar peptidoglycan of rod-shaped cells has traditionally been considered to be “inert” such that it experiences reduced rates of remodeling relative to the length of the cell. What makes poles behave as though they are inert, however, is unknown. Here, we use a dye that stains peptidoglycan either during synthesis or remodeling to show that in the absence of Min, polar peptidoglycan is indistinguishable from the rest of the cell. How MinD, a protein in the cytoplasm would inhibit peptidoglycan remodeling extracellularly is unclear, but we note the *B. subtilis* Min system interacts with a polarly-localized multi-pass transmembrane protein, MinJ (43, 44). MinJ has recently been shown to interact with RodZ, a protein involved in peptidoglycan synthesis/remodeling, and perhaps the MinCDJ complex keeps RodZ away from the poles (84–87). Moreover, why mincells fail to grow is poorly understood, but perhaps, they quickly lose the ability to remodel their peptidoglycan. The loss of remodeling capacity could be due to both the degradation of a single required protein and its corresponding transcript, but we note that the most substantial difference between mother cells and minicells of a *minD* mutant is the absence of the chromosome. Thus, the chromosome may not only direct the cell division cycle, but its presence may either directly or indirectly dictate cell envelope remodeling, and we note that RodZ is a transmembrane protein with a cytoplasmic DNA binding domain (58, 84, 86, 88).

## ACKNOWLEDGEMENTS

We thank Yves Brun and Mike VanNieuwenhze for material support and helpful comments. The work was supported by NIH grant R35 GM131783 to DBK and NIH grant R01 GM113172 to SJ.

## FIGURE LEGENDS

**Figure S1. Diagram of the microfluidic device.** Black lines indicate microfluidic channels. Open circles indicate media/reagent input/output reservoirs. Closed circles indicate the location of peristaltic valves. Three valves in parallel indicate a peristaltic pump. Location of the nanochannel array imaged by fluorescence microscopy indicated by a red box. The same device was used previously in (Baker et al., 2016).

**Figure S2.** Data histograms of FtsZ parameters used to generate Figure 4. 100 cells were manually tracked and measured through a cell cycle to generate the data for each panel. Data are presented as gray bars for wild type (DK5133) and as blue bars for the *minD* mutant (DK5155). A) Frequency histogram of the Z-ring appearance period defined as the time between the formation of one Z-ring and the next Z-ring. B) Frequency histogram of the Z-ring persistence period defined by the time between the formation of a Z-ring and the disappearance of that Z-ring. Note, no data are provided for the *minD* mutant as the Z-rings of a *minD* mutant did not disappear and the persistence period was effectively infinite. C) Frequency histogram of the Z-ring polar duration defined as the time between a septation event and the disappearance of the Z-ring resulting from that septation events. Note, no data are provided for the *minD* mutant as the Z-rings of a *minD* mutant did not disappear and the polar duration period was effectively infinite. D) Frequency histogram of the Z-ring medial delay defined as the time between a septation event and the formation of a Z-ring that will eventually give rise to the next medial division event. Note that the Z-ring medial delay of the *minD* mutant was often negative because the medial Z-ring that would eventually promote cell division was formed in the preceding generation. E) Frequency histogram of the Z-ring maturation period defined as the time between Z-ring formation and when that Z-ring achieves peak local intensity. F) Frequency histogram of the cytokinetic period defined as the time between Z-ring formation and septation directed by that Z-ring.

**Figure S3. The *minD* mutant accumulates and loses fluorescent BADA signal more rapidly than the wild type.** A) Total BADA fluorescence intensity after a 4-min staining impulse and washout of the BADA stain in the wild type (DK4393). Points are the average, and whiskers are the standard deviation of over 500 measurements. B) Total BADA fluorescence intensity after a 4-minute staining impulse and washout of the BADA stain in the *minD* mutant (DK4407). Points are the average, and whiskers are the standard deviation of over 500 measurements. Cyan indicates measurements of the long mother cells with nucleoids, and magenta indicates measurements of minicells.

**Figure S4. A minicell redistributed peptidoglycan stain from its old cell pole to its new cell pole.** Cells were stained with BADA for 4 min, then media lacking BADA was introduced to wash out the dye, and a kymograph of one channel was generated. Here, we highlight an instance in which a minicell was formed by polar division soon after the dye had been removed such that old pole was stained but the division plane and resulting new pole was not (white caret). Over time, the fluorescent signal incorporated into the peptidoglycan becomes uniformly redistributed around the circumference of the minicell (black caret).

**Movie S1: Wild type growth in microfluidic channels.** Constitutive cytoplasmic mCherry, false colored red. Strain DK5133. Movies are 200 frames over 400 min at a rate of 1 frame/2 min.

**Movie S2: Wild type growth in microfluidic channels with fluorescent FtsZ.** Constitutive cytoplasmic mCherry, false colored red, and a mNeongreen-FtsZ, false colored green. Strain DK5133. Movies are 200 frames over 400 min at a rate of 1 frame/2 min.

**Movie S3: *minD* mutant growth in microfluidic channels.** Constitutive cytoplasmic mCherry, false colored red. Strain DK5155. Movies are 200 frames over 400 min at a rate of 1 frame/2 min.

**Movie S4: *minD* mutant growth in microfluidic channels with fluorescent FtsZ.** Constitutive cytoplasmic mCherry, false colored red, and a mNeongreen-FtsZ, false colored green. Strain DK5155. Movies are 200 frames over 400 min at a rate of 1 frame/2 min.

**Movie S5: Wild type growth in microfluidic channels with fluorescent BADA that stains newly synthesized/remodeled peptidoglycan.** Constitutive cytoplasmic mCherry, false colored red, and stained with a 4 minute pulse of BADA, false colored green. Strain DK4393. BADA was washed out of the channel for 4 minutes after which, imaging was recommenced. Movies are 60 frames over 120 min at a rate of 1 frame/2 min.

**Movie S6: *minD* growth in microfluidic channels with fluorescent BADA that stains newly synthesized/remodeled peptidoglycan.** Constitutive cytoplasmic mCherry, false colored red, and stained with a 4 minute pulse of BADA, false colored green. Strain DK4407. BADA was washed out of the channel for 4 minutes after which, imaging was recommenced. Movies are 60 frames over 120 min at a rate of 1 frame/2 min.

